# U1 snRNP regulates cancer cell migration and invasion

**DOI:** 10.1101/730515

**Authors:** Jung-Min Oh, Christopher C. Venters, Chao Di, Anna Maria Pinto, Lili Wan, Ihab Younis, Zhiqiang Cai, Chie Arai, Byung Ran So, Gideon Dreyfuss

**Affiliations:** Howard Hughes Medical Institute, Department of Biochemistry and Biophysics, University of Pennsylvania School of Medicine, Philadelphia, PA 19104-6148

## Abstract

Stimulated cells and cancer cells have widespread shortening of mRNA 3’-utranslated regions (3’UTRs) and switches to shorter mRNA isoforms due to usage of more proximal polyadenylation signals (PASs) in the last exon and in introns. U1 snRNA (U1), vertebrates’ most abundant non-coding (spliceosomal) small nuclear RNA, silences proximal PASs and its inhibition with antisense morpholino oligonucleotides (U1 AMO) triggers widespread mRNA shortening. Here we show that U1 AMO also modulates cancer cells’ phenotype, dose-dependently increasing migration and invasion in vitro by up to 500%, whereas U1 over-expression has the opposite effect. In addition to 3’UTR length, numerous transcriptome changes that could contribute to this phenotype are observed, including alternative splicing, and mRNA expression levels of proto-oncogenes and tumor suppressors. These findings reveal an unexpected link between U1 regulation and oncogenic and activated cell states, and suggest U1 as a potential target for their modulation.

## Introduction

Widespread shortening of mRNA 3’-utranslated regions (3’UTRs) and switches to short mRNA isoforms is a common feature and contributing factor to cell stimulation, seen in immune cells and neurons, and oncogenicity^1-6^. These shortening events occur due to a shift in usage of more upstream polyadenylation signals (PASs) in the last exon and in introns. In quiescent cells, these PASs are generally silenced by U1 snRNP (U1), vertebrates’ most abundant non-coding small nuclear RNP, which is necessary for production of full-length RNA polymerase II transcripts from protein-coding genes and long noncoding RNAs^7^. This U1 activity, called telescripting, requires RNA base-pairing of U1 snRNA 5’-end, which is also required for U1’s role in 5’ splice site (5’ss) recognition. U1 antisense morpholino oligonucleotides (U1 AMO), which inhibits U1:pre-mRNA base-pairing, triggers widespread premature cleavage and polyadenylation (PCPA) as well as inhibits splicing^8,9^. Transfection of a high U1 AMO dose that masks all, or nearly all, of U1 snRNA 5’-end causes drastic PCPA from cryptic PASs frequently found in the 5’-side of the intron in pre-mRNAs of thousands of genes^8,9^, especially in long introns of large genes (>39kb). In contrast, small genes (<6.8kb) are generally PCPA-resistant and many of them are up-regulated in this environment. Importantly, these small genes are enriched in functions related to cell survival and acute cell stress response^7^. The drastic PCPA from high U1 AMO in many genes obscures other effects that are more readily detected with low U1 AMO doses. These changes include 3’UTR shortening (shifts to usage of more proximal PASs in tandem PASs) and the production of shorter mRNA isoforms from PAS usage in introns^9^. This revealed that low U1 AMO recapitulates known shifts to shorter mRNA isoforms that occurs in stimulated neuronal cells. We have shown previously that the stimulation-induced rapid transcription up-regulation creates transient telescripting deficit because U1 levels cannot rise in step with the transcription surge from immediate early genes^9^. U1 synthesis is a slower process involving nuclear export of pre-snRNAs, SMN complex-mediated snRNP assembly in the cytoplasm, and re-import to the nucleus^10^. Consequently, some pre-mRNAs transcribed in this time window (∼2-6 hours post stimulation) are processed to shorter mRNA isoforms due to PCPA in introns. For example, pre-mRNA processing of *homer-1*, which encodes a synaptogenesis scaffold protein, shifted from full-length mRNA to a shorter mRNA isoform that encodes antagonistic activity due to PCPA in an intron^11^. Importantly, low U1 AMO recapitulated the same isoform switching^9^. Here, we investigated if low U1 AMO could also modulate cell phenotype.

## Results and Discussion

### U1 level changes modulate hallmark phenotypes of cancer cells in vitro

We used standard *in vitro* assays to determine if moderate U1 inhibition has an effect on proliferation, migration, or invasion of cancer cells, which serve as quantitative measures of oncogenic phenotype. Various low U1 AMO doses (2.5 ∼ 250 pmole) or control, non-targeting AMO (cAMO) were transfected into HeLa cells, a cervical carcinoma cell line (Fig. 1). These U1 AMO doses masked ∼15-30% of U1 snRNA 5’-ends, making it inaccessible for base-pairing corresponding to the U1 AMO dose^8,9^. As shown in Fig. 1a, 62.5 pmole U1 AMO moderately increased (38%) cell proliferation after 48-72 hours. Higher U1 AMO doses (≥250 pmole) were toxic and reduced overall cell numbers. Remarkably, low U1 AMO dose-dependently enhanced cell migration and invasion by up to ∼500% in 24 hours with peak activity at 62.5 pmole (Fig. 1b-e). The increased migration and invasion reflect true enhancements that could not be accounted for by the comparatively small increase in cell number (Fig. 1a). U2 AMO, which interferes with U2 snRNP’s function in splicing^12^, did not enhance any of these phenotypes over the entire dose range, indicating that the specificity of the U1 AMO effects (Supplementary Fig. 1).

**Fig 1.**
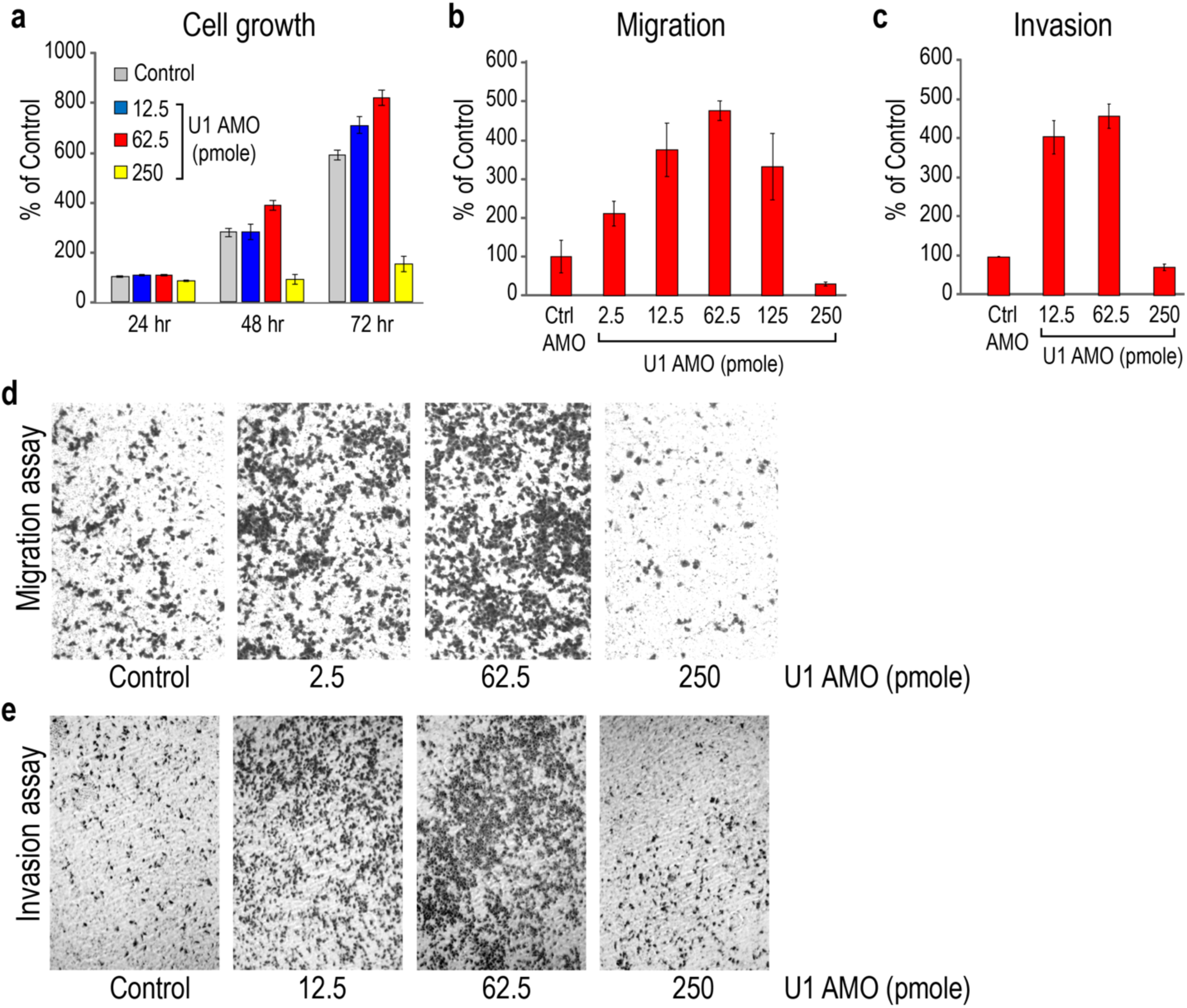
Low U1 AMO dose-dependently enhances migration and invasion of HeLa cells *in vitro*. **a** HeLa cells were transfected with the indicated amounts of U1 or control AMOs. Cell proliferation was determined from cell numbers using the Cell Titer-Glo Luminescent Cell Viability Assay at the times indicated. Data are represented as mean ± standard deviation (SD). **b-c** Migrating and invading cells were measured 24 hours post-transfection. 1.5 x 10^5^ and 2.5 x 10^5^ cells were cultured for the migration and invasion assays, respectively. Data are represented as mean ± SD. (**d**,**e**) Migrating and invading cells were stained 24 hours after transfection and visualized by phase-contrast microscopy (10X magnification).

To further examine the effects of U1 on cell phenotype, we over-expressed U1 (U1 OE) from a plasmid carrying U1 snRNA gene with its native promoter and termination elements. This achieved 20% and 40% U1 level increases compared to empty plasmid by transfecting 1µg and 1.5µg of this plasmid, respectively (Supplementary Fig. 2). The U1 OE significantly and dose-dependently attenuated cell migration (25-50%) and cell invasion (25-65%) after 24 hours (Fig. 2). U1 OE dose-dependently increased the amount of U1 snRNP, determined by the amount of U1 snRNA in anti-Sm immunoprecipitations (data not shown).

**Fig 2.**
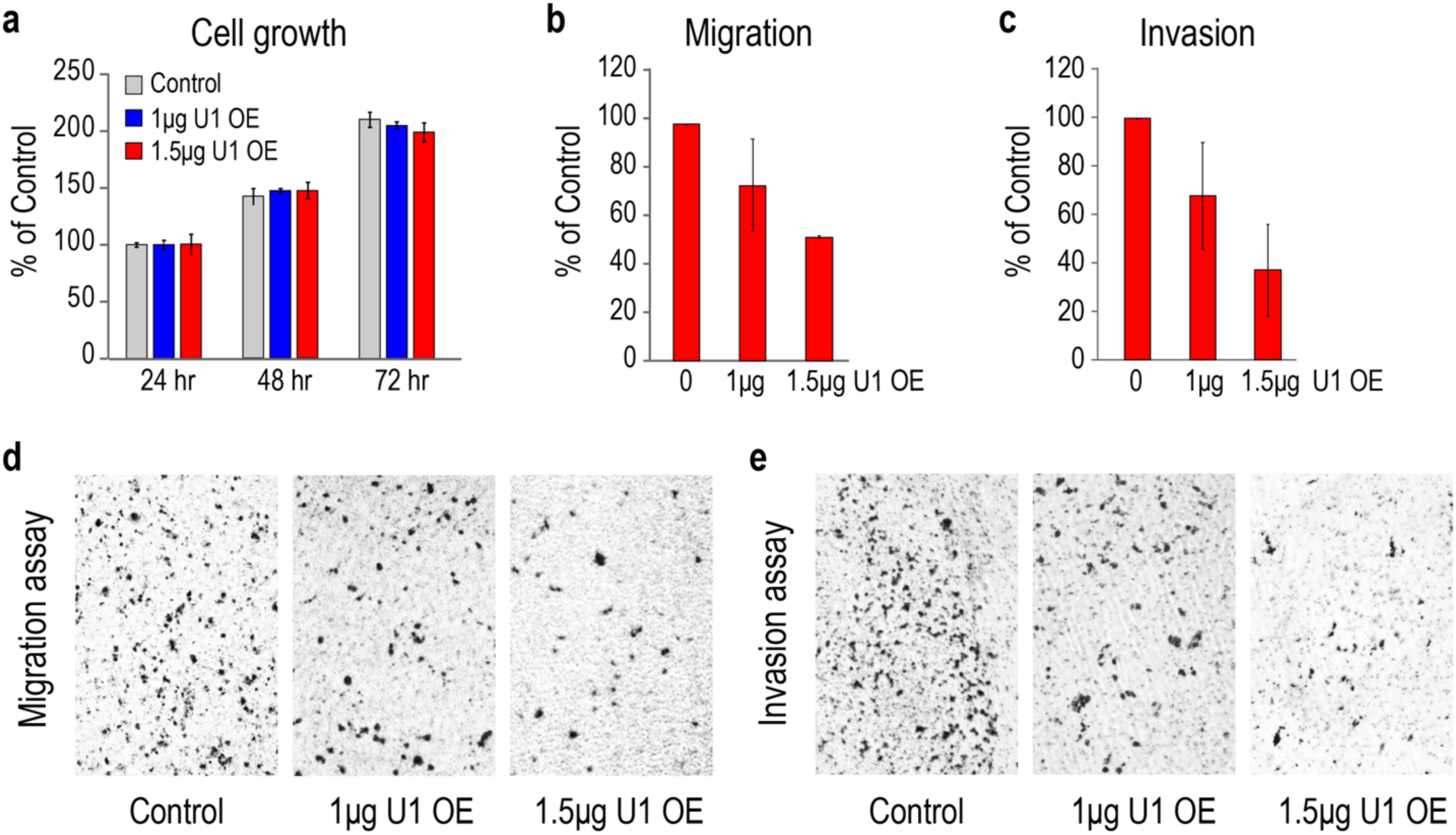
U1 over-expression decreases oncogenicity of HeLa cells *in vitro*. HeLa cells were transfected with control empty vector or U1 expression plasmids. **a** Cell growth determined in 24 hours intervals. The data are represented as mean ± SD. **b-c** Cell migration and invasion were measured 24 hours after transfection. The bars indicate mean ± SD. **d-e** Cell migration and invasion were determined 24 hours post-transfection and imaged by phase-contrast microscopy (10X magnification).

Similar experiments on other cancer cells, including human lung adenocarcinoma (A549) and breast adenocarcinomas (MCF-7 and MB-231), demonstrated the generality of the U1-related effects on phenotype. U1 AMO enhanced migration of A549, MCF-7 and MB-231 by 58-72% and increased invasion of A549 and MB-231 by 53-64% (Supplementary Table 1). Conversely, U1 OE attenuated the migration and invasion of these cell lines by ∼50% compared to the control levels (Supplementary Table 1).

### U1 level changes cause numerous and diverse transcriptome changes

We used high-throughput RNA sequencing (RNA-seq) to determine transcriptome changes resulting from U1 level modulation. To enhance detection of nascent RNAs, as U1 functions co-transcriptionally^7^,we metabolically labeled RNAs with 4-thiouridine (4-shU) for two hours (at 6-8 hours post-transfection with AMOs and 22-24 hours of U1 OE), and sequenced the thiol-selected RNAs. Reads were mapped to the human genome (UCSC, hg38) and filtered for unique alignments. The statistics of the RNA-seq datasets, normalized for sequencing depth as reads per million (RPM), are shown in Supplementary Table 2. Over 9,100 genes expressed to RPKM > 1 in the control samples for U1 AMO and U1 OE, and were included in further analysis. This revealed numerous and diverse transcriptome changes caused by U1 AMO, including in mRNA expression, 3’UTR length, and alternative splicing affecting thousands of genes. The number of genes affected by each type of change are listed in Supplementary Table 3. Frequently, more than one type of change was detected in transcripts of the same gene (18-47%; Supplementary Fig. 3). Generally, the number of events for each type of change increased with U1 AMO dose and there was extensive overlap (31-97%) between events detected at the two doses tested (Supplementary Table 3).

Confirming earlier observations from datasets with lower resolution and sequencing depth^9^, low U1 AMO elicited widespread 3’UTR shortening readily detected in genome browser views, for example, *TOP2A, TFRC* and *CDC25A*. (Fig. 3). A common approach to identify changes in locations of 3’-poly(A)s is to use oligo(dT)-primed RNA-seq^13^. However, it is not quantitative and does not provide adequate information on overall transcriptome changes due to 3’ bias. An alternative method, DaPars, uses a regression model to deduce alternative PAS usage among multiple tandem 3’UTR PASs from standard RNA-seq^14^. Frequently, 3’UTRs have multiple tandem PASs, resulting in complex mixtures of mRNAs with various 3’UTR lengths that make it difficult to resolve by DaPars. We reasoned that a shift in usage to proximal 3’UTR PASs would decrease the overall amount of transcription in the 3’UTR, while a shift to more distal PASs will increase it. As changes in 3’UTR amount could result from mRNA expression level changes alone, we calculated the ratio of RNA-seq reads in the 3’UTR to the reads in the portion of the CDS in the same last exon (heretofore LECDS). This normalizes 3’UTR signal to mRNA expression. A decrease or increase in LECDS in a sample compared to control suggests 3’UTR shortening or lengthening, respectively.

**Fig 3.**
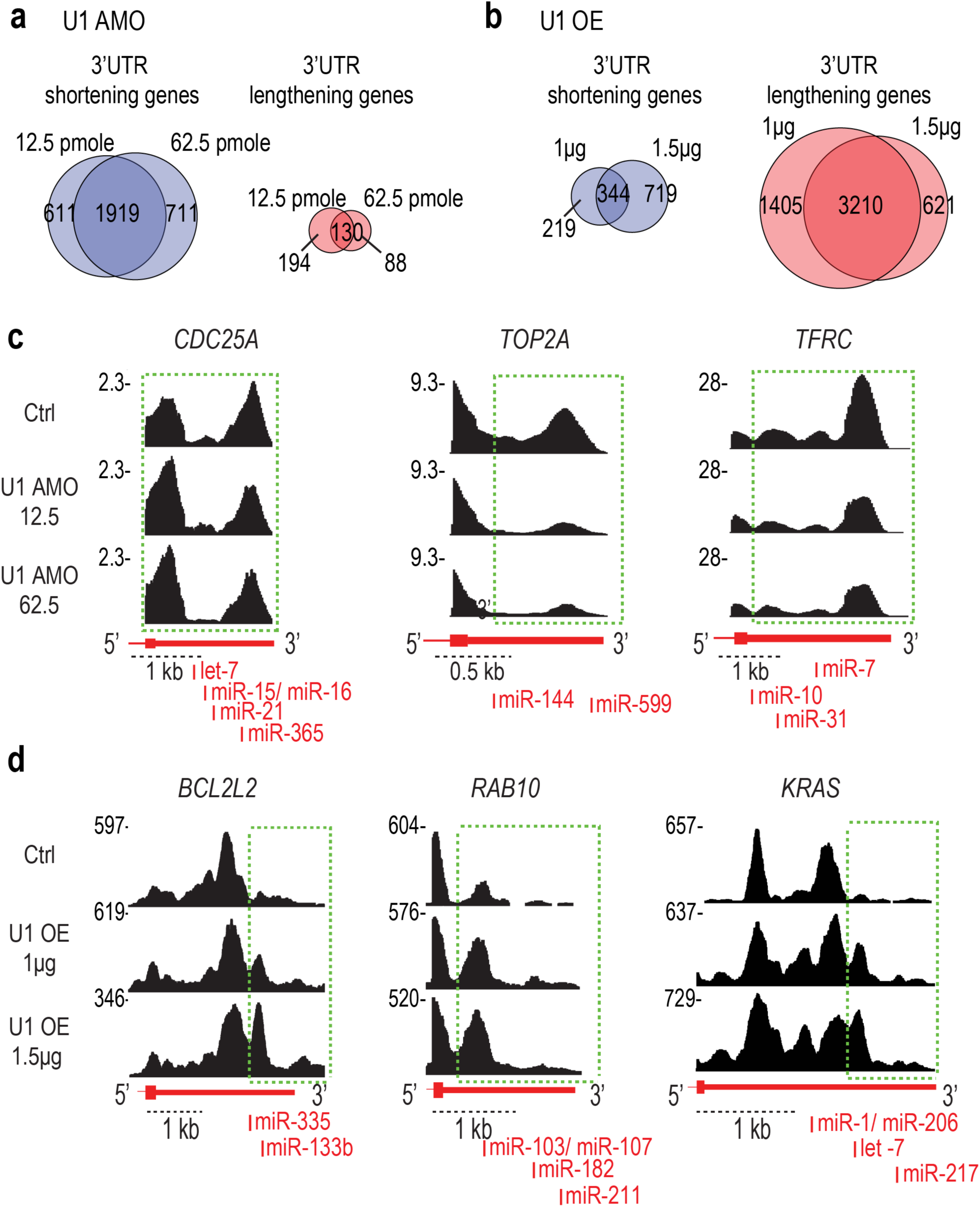
Examples of genes with 3’UTR shortening by moderate U1 decrease (inhibition with U1 AMO) and genes with 3’UTR lengthening from U1 over-expression. Venn diagrams of LECDS-identified genes affected by 3’UTR shortening versus 3’UTR lengthening in U1 AMO 12.5 and 62.5 pmole (**a**) and U1 OE 1 and 1.5 µg (**b**) samples. **c** Cells transfected with control or U1 AMO (12.5 and 62.5 pmole, 8 hours) were labeled with 4-thiouridine to select and sequence nascent transcripts. RNA-seq maps of representative genes from two biological experiments are shown. Green dotted boxes indicate the affected regions where shortening in the 3’UTR occurs. **d** Cells transfected with control or U1 expression plasmids (1 µg and 1.5 µg, 24 hours) were labeled with 4-thiouridine to select nascent transcripts. Green dotted boxes indicate the regions where there was a switch towards longer 3’UTRs. miRNA target sites that are involved in cancer and expressed in HeLa are shown.

LECDS identified 3’UTR shortening in 1,919 genes in both 12.5 and 62.5 U1 AMO (p ≤ 0.01), and only a small fraction (130 genes) had more reads in 3’UTRs (Fig. 3a). In contrast, U1 OE caused a 3’ increase in reads (longer 3’UTRs) in 3,210 mRNAs, while it shortened 3’UTRs in only 344 genes (Fig. 3b). The large overlap of the lengthened (∼84%) and shortened genes (∼76%) in both U1 OE (1µg and 1.5µg) and U1 AMO samples (U1 AMO 12.5 and 62.5), respectively, showed a strong validation of the RNA-seq data from separate biological experiments. Comparison of the 3’UTR changes identified using LECDS to those identified using DaPars showed an overlap of 79% in the samples. Moreover, LECDS identified 2,630 more events than DaPars (Supplementary Fig. 4), which is likely due to increase usage at multiple PASs rather than a strong increase in usage of a single alternative PAS. Further confirmation of 3’UTR shortening and lengthening is shown for select examples in Fig. 3c-d, and 3’RACE validation data are presented in Supplementary Fig. 5. Notably, the fraction of genes that had 3’UTR length changes corresponded to the U1 AMO or OE dose, and U1 OE caused transcription in some genes to extend farther downstream from canonical gene ends. Thus, U1 has a role in regulation of both proximal and distal PASs.

As expected for U1’s role in splicing, U1 AMO caused alternative splicing (AS) changes in ∼700 genes, including A5’SS and A3’SS, cassette exon (CE) and intron retention (IR) events (Supplementary Table 3 and Fig. 4)^15^. Examples of AS and confirmation of select cases by RT-PCR are shown in Fig. 4b and Supplementary Fig. 6. Many of the observed AS changes could contribute to the oncogenic phenotype. For example, ataxia-telangiectasia, a cancer predisposition disorder caused by mutations in the ATM gene^16^, has two introns retained (intron 1 and 33) and is down-regulated in U1 AMO compared to cAMO (Figs. 4b and 5).

**Fig 4.**
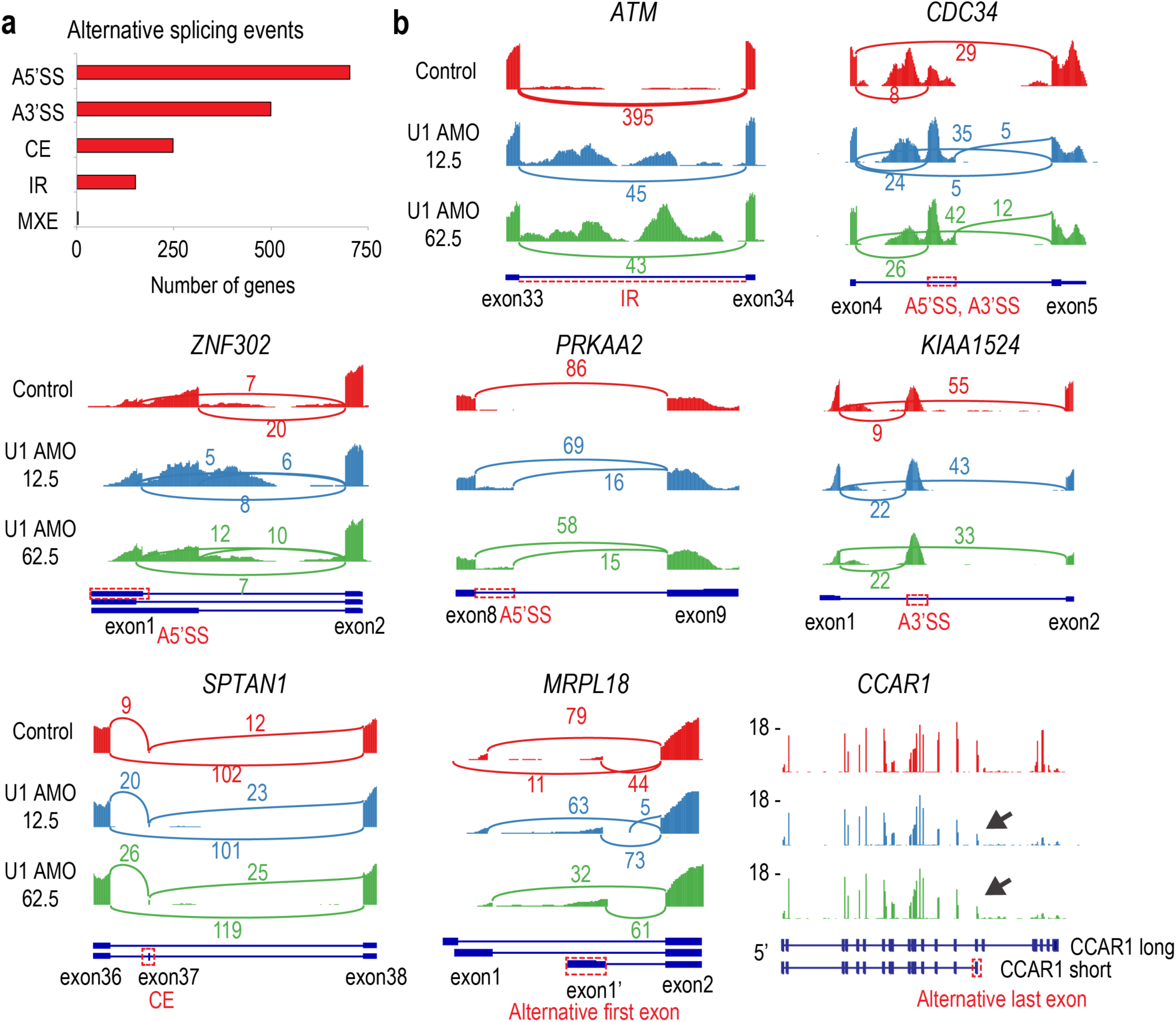
Low U1 AMO causes widespread alternative splicing changes in HeLa cells. **a** Summary of alternative splicing events as identified with JUM analysis. Each alternative splicing event happened in both low U1 AMO 12.5 and 62.5 pmole in HeLa cells. (A5’SS: alternative 5’ splice site; A3’SS: alternative 3’ splice site; CE: cassette exon; IR: intron retention; MXE: mutually exclusive exons) **b** Sashimi plots and genome browser view of each alternative splicing change in the low U1 AMO 12.5 and 62.5 pmole.

**Fig 5.**
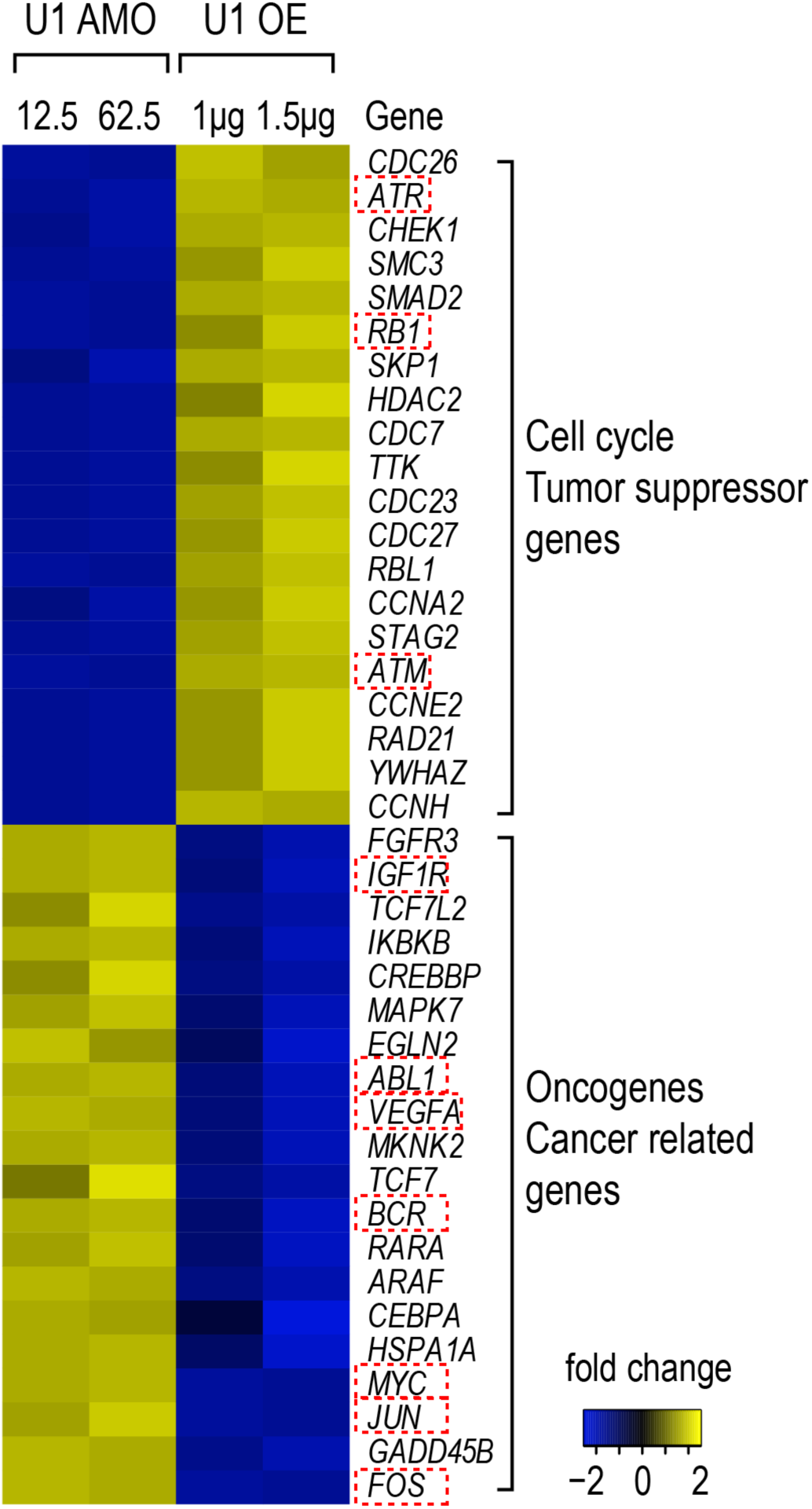
U1 AMO up-regulated oncogenes and down-regulates tumor suppressor genes and U1 overexpression antagonizes the effect in HeLa cells. Heat map showing genes related to cell cycle and tumor suppressor function are down-regulated in U1 AMO and up-regulated in U1 OE. Oncogene and cancer related genes are up-regulated in U1 AMO and down-regulated in U1 OE (1.5-fold change, p-value < 0.01 by GFold^53^).

Gene Ontology analysis (GO term) revealed expression level changes induced by U1 level modulation in multiple oncogenes, tumor suppressors and cell cycle related genes. For example, *ATR, RB1, ATM* and several *CDC* (Cell Division Cycle) genes were down-regulated in U1 AMO, but up-regulated in U1 OE. On the other hand, oncogene and cancer related genes, such as *MYC, FOS, JUN*, and growth promoting genes were up-regulated in U1 AMO, but down-regulated in U1 OE (Fig. 5 and Supplementary Fig. 7). The expression down-regulations at 6 hours post-U1 AMO transfection could also be caused by PCPA or by splicing changes, for example by intron retention that introduces a premature translation stop codon and triggers nonsense mediated mRNA decay. Many of the transcriptome changes could also be due to secondary cellular effects, especially at the longer time point (24 hours), for example due to changes in the level of other transcription and splicing factors. Nevertheless, concomitant up-regulation of oncogenes and down-regulation of tumor suppressor genes likely contribute to the phenotypic changes in the wake of U1 AMO. Among genes affected by U1 AMO are splicing factors, which have been linked to myelodysplastic syndromes, chronic lymphocytic leukemia and other cancers (Supplementary Fig. 8)^17-20^.

### U1 level changes alters expression of cancer genes

To explore the potential role of U1 dependent 3’UTR length changes on the oncogenic phenotype we observed, we interrogated our gene set against cancer gene databases [Sanger^21^ and UCSF (waldman.ucsf.edu/GENES/completechroms.html)]. These archives include oncogenes induced through various mechanisms, including mutation, chromosomal translocation, or loss of miRNA repression, and the analysis revealed that a large number of oncogenes incurred 3’UTR length changes. The oncogenes affected at each U1 level, 204 in total, are listed in Supplementary Table 4 and include cell cycle regulation (*CDC25A, CCNB1*), apoptosis (*BCL6, BRCA1*), cell migration (*FGFR1, FYN*), extracellular matrix remodeling (*TIMP2*), signaling (*EGFR*), transcription (*WNT5A*), metastasis and tumor progression (*EWSR1, APC, BRAF*).

There are numerous examples of cancers resulting from oncogene up-regulation due to loss of miRNA repression, either because the relevant miRNA is down-regulated or its target in the 3’UTR has been removed^22-25^. U1 level changes recapitulated the same miRNA 3’UTR target elimination or restoration in many genes (Fig. 3c-d). For CDC25A, an essential phosphatase for the G1-S transition, 3’UTR shortening eliminates several miRNA-binding sites including let-7, miR-15 and miR-21 (Fig. 3c). An increase in CDC25A protein due to alleviation of miRNA-mediated repression exacerbates hepatic cyst formation and colon cancer^26-28^. 3’UTR shortening in RAS oncogene family member, *RAB10*, which removes target sites for miRs-103/107, is found in numerous cancer cell lines and results in a dramatic increase of this protein^4^. As shown in Fig. 3d, 3’UTR of *RAB10* is already shortened in HeLa cells, and U1 OE reverses this shortening and restores the corresponding miRNA binding sites. Similarly, U1 OE lengthened 3’UTR of *KRAS* to include a let-7 binding site (Fig. 3d). This region contains a polymorphism that has been shown to impair let-7 binding, and is prognostic for breast cancer aggressiveness^29^. U1 AMO also did not change the associated miRNA expression (Supplementary Fig. 9). In addition to loss or gain of miRNA-binding sites, 3’UTR length or amount changes can affect other mRNA regulating elements, such as AU-rich elements (AREs), with roles in cancer phenotype. For example, U1 OE-induced 3’UTR lengthening in *c-Fos*, a gene controlling cell proliferation, differentiation, survival and tumorigenesis^30,31^, restores an ARE and strongly decreases mRNA level (>2-fold, data not shown).

Several other factors that can regulate 3’UTR length have been described, particularly components of the cleavage and polyadenylation machinery (CPA)^13,32-36^. Knockdown of CPSF6/CFIm68, CPSF5/CFIm25, and PABPN1 cause widespread 3’UTR shortening^35,37,38^. Furthermore, CPSF5 and CPSF6 are down-regulated in cancers and their knockdowns in cell lines enhances oncogenicity^14,35,39,40^. It is possible that these CPA factors work in concert with U1. 3’ UTR shortening analysis of the RNA-seq of CPSF5 knockdown in HeLa cells by LECDS identified 1,210 genes (63% overlap with U1 AMO) that are also shortened in U1 AMO (Supplemental Fig. 10), suggesting potentially extensive commonality of targets. A recent study described that U1 over-expression in PC-12 rat neuronal cell line up-regulated cancer related genes, including MYC and FOS^41^. While these results appear to be inconsistent with our observations, they may be explained by the different cell types used in these studies and by different U1 levels relative to transcription.

The large number and diversity of transcriptome changes make it impossible to attribute the phenotypic change to specific transcriptome changes. However, many of the changes could be oncogenic by themselves. Thus, a surprising overall observation is that at normal U1 and transcription levels, U1 snRNP has a tumor suppressor-like function. U1’s ability to regulate such genes, and loss of these functions as less U1 is available, fulfills the definition of (an overarching) tumor suppressor^42,43^. Mutations in several splicing factors, particularly U2-associated factors and SR proteins, have been shown to drive many cancers^17,18^. However, the potential role of U1 in cancer has not been tested previously. Our findings demonstrate that U1 homeostasis is necessary for maintaining normal expression of many of the same factors, as well as numerous other cancer-related genes. In this vein, it is intriguing that the major U1 gene cluster in humans at chr1p36.13 flanks a breakpoint linked to many cancers^44^. The potential effect of these genomic abnormalities on U1 levels remains to be determined. Mounting evidence indicate that U1 AMO and U1 over-expression mimic biological processes^7,9,45-48^. Thus, these experimental tools for dose-dependent modulation of U1 should facilitate studies on cancer and stimulated cells behavior, and suggest U1 as a potential clinical target for a wide range of disorders.

## Methods

### Cell culture and treatment

Cells (HeLa, A549, MCF-7 and MB-231) were maintained in DMEM supplemented with 10% fetal bovine serum (FBS), 10 units/ml penicillin and 10 μg/ml streptomycin at 37°C and 5% CO_2_. U1, U2 antisense and control morpholino oligonucleotide sequences were as previously described^8^. Oligonucleotides were transfected into 1 x 10^6^ cells using a Neon transfection system (Invitrogen) according to the manufacturer’s instruction to achieve the desired final doses indicated in the text and figures.

### Proliferation assay

The Cell Titer-Glo Luminescent Cell Viability Assay (Promega) was used according to the manufacturer’s instructions to measure cell proliferation. The cells were transfected with AMOs and seeded in triplicate in 96-well plates at a density of 1 x 10^4^ cells per well. Cells were incubated in media containing 1% FBS for 3 days and proliferation was measured every 24 hours.

### Migration and invasion assays

For migration studies, a standard assay was used to determine the number of cells that traversed a porous polycarbonate membrane in response to a chemo-attractant (higher serum concentration)^49,50^ using the Cytoselect 24-well cell migration assay (Cell Biolabs). Invasion was measured using BD BioCoat Matrigel invasion chambers (BD Bioscience). In both assays, cells were transfected with AMOs, and 1.5 x 10^5^ and 2.5 x 10^5^ cells per well were seeded in an upper chamber in serum free media for the migration and invasion assay, respectively. The lower chamber was filled with media containing 10% FBS. After 24 hours, cells passing through polycarbonate membrane were stained and counted according to the manufacturer’s instructions.

### Metabolic RNA labeling, isolation and RNA-seq

4-thiouridine (250 μM) was added to cells between 6-8 hours after U1 AMOs transfection. Total RNA was extracted with Trizol (Invitrogen) and poly(A) RNA purified on Oligotex beads (Qiagen). Free thiols on poly(A) mRNA were reacted with 0.2 mg/ml biotin-HPDP for 2 hours to label RNA that incorporated 4-thiouridine. RNA was then purified on M-280 streptavidin Dynabeads (Invitrogen), cDNA was synthesized using Ovation RNA-Seq System V2 (NuGEN) and libraries for Illumina sequencing were constructed using Encore NGS Library System (NuGEN) according to the manufacturer’s instructions.

### Mapping RNA-seq reads

RNA-seq reads were aligned to reference genome UCSC/hg38 using STAR^51^ with default settings. Reads per exon were grouped, from which RPKM (Read Per Kilobase of exon model per Million mapped reads)^52^ values were calculated. Only genes with RPKM ≥ 1 were included in further analyses.

### Differential gene expression, alternative splicing and miRNA analyses

The GFold algorithm^53^ was used to calculate differential expression using the default parameters for detecting reliable expression change on all exons of the longest full-length isoform excluding the 3’UTR. A 1.5-fold change was set with a p-value < 0.01 to identify genes with significant expression change. The Junction Usage Model (JUM)^15^ analysis was performed to identify differentially spliced isoforms in experimental samples compared to controls and quantify their expression levels by computing the ΔPSI (difference of Percent Spliced Isoform). Significant differential alternative splicing events were detected by performing a χ^2^ likelihood-ratio test followed by Benjamini–Hochberg (BH) multiple testing with an adjusted P-value ≤ 0.01. For stringent inclusion, we also restricted ΔPSI >=10%. TargetScan miRNA regulatory sites were downloaded from UCSC Genome browser (http://genome.ucsc.edu/). The coordinate of each predicted miRNA site was compared to those genes for which the 3’UTR showed changes in length. For miRNA microarray, HeLa cells were transfected with control or 62.5 pmole U1 AMO for 8 hours. Total RNA was extracted using miRNeasy kit (Qiagen) according to manufacturer recommendations. GeneChip miRNA 3.0 array (Affymetrix) was used to determine the miRNA expression and experiments and analysis were performed by Molecular Profiling Facility at University of Pennsylvania.

### LECDS calculations

We used both the 3’UTR and coding sequence of the terminal exon to determine the change in reads in 3’UTR of the longest (full-length) isoform. As 3’UTR reads can change with expression level changes, such as transcription up- or down-regulation, 3’UTR signals were compared to that from the last exon’s coding sequence (LECDS). If the 3’UTR reads are significantly decreased or increased relative to those from the LECDS, the 3’UTR is called as shortening or lengthening, respectively. The significance of this read change was detected using a Fisher’s Exact test followed by Benjamini–Hochberg (BH) multiple testing with an adjusted P value ≤ 0.01. This provides an accurate read-out of the net change in 3’UTR expression in a gene’s transcript and can be used based on total RNA-seq without specialized poly(A) mapping.

## Acknowledgements

We thank members of our laboratory for helpful discussions and comments on the manuscript. This work was supported by the US National Institutes of Health (R01GM112923 to G.D.). G.D. is an Investigator of the Howard Hughes Medical Institute.

## Author contributions

JM, AMP, LW, IY and GD conceived and designed the project. JM, AMP, LW, IY, CA, ZC, and BRS performed the experiments. CCV and CD performed the informatics analyses. JM, CCV, CD and GD wrote the manuscript with input from all the authors. GD was responsible for the project’s performance.

## Data availability

All sequencing data described are available on GEO under the accession number (to be assigned).

## Code availability

Code for the analyses described in this study is available from the corresponding author upon request.

## Supplementary Figures and Tables

**Supplementary Fig 1.**
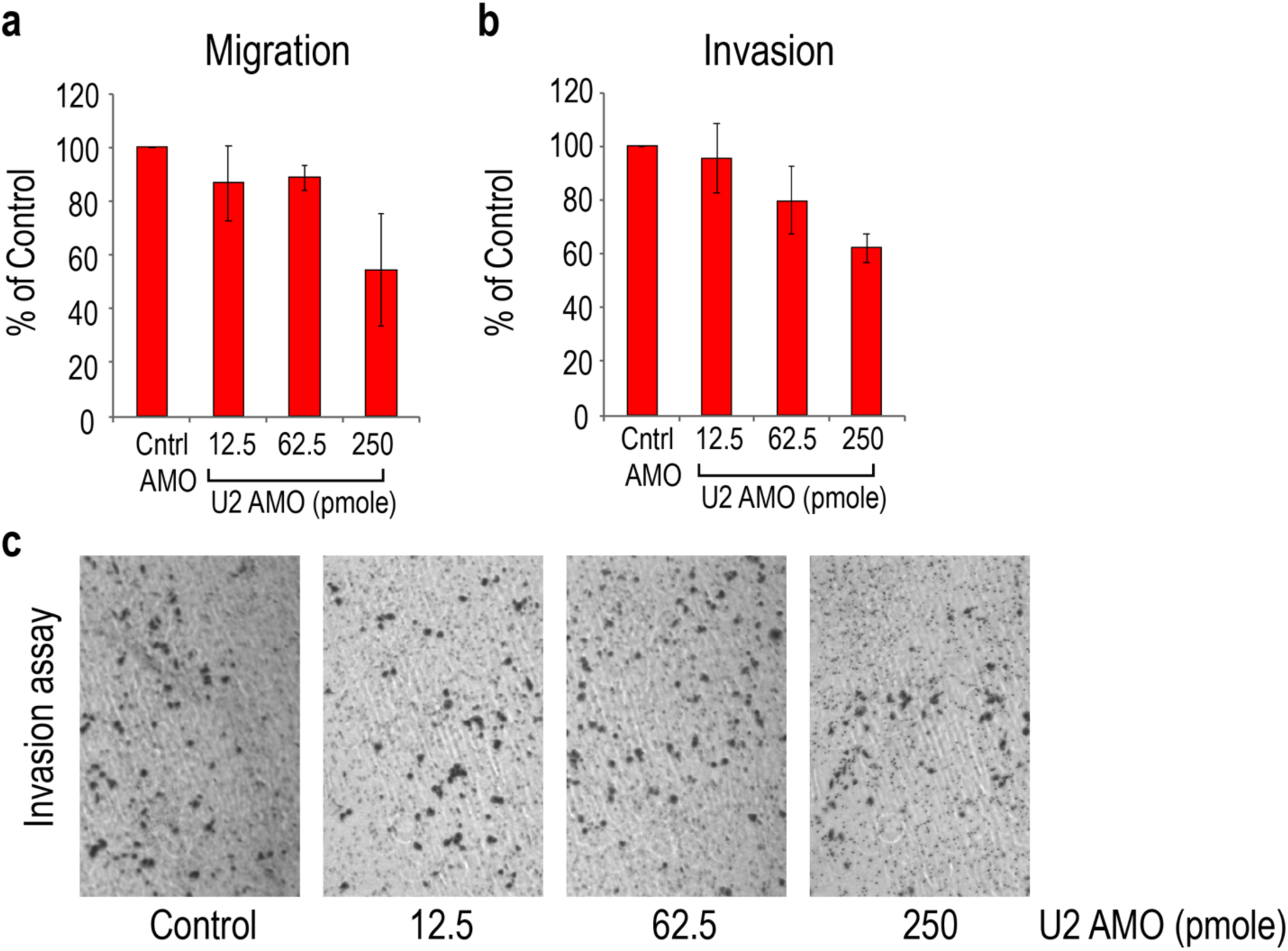
U2 AMO does not enhance migration and invasion of HeLa cells *in vitro*. **a** Migration assay (**b**) invasion assay, and (**c**) phase-contrast imaging 24 hours following transfection with control or U2 AMO (12.5-250 pmole) were performed as described in Fig 1. Data are represented as mean ± SD. U2 AMO at 250 pmole decreased cell numbers.

**Supplementary Fig 2.**
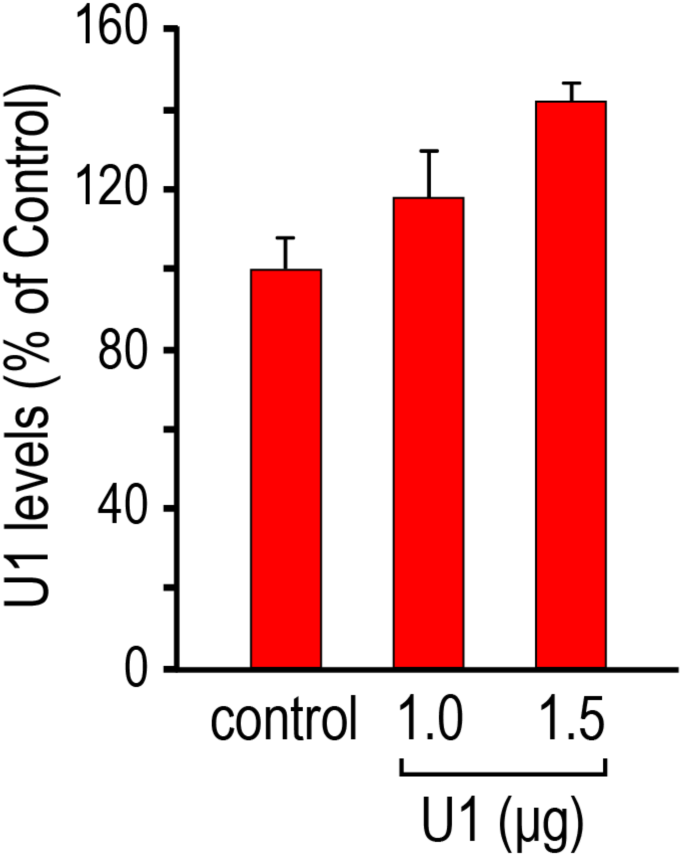
Expression of exogenous U1 snRNA. HeLa cells were transfected with increasing concentrations of a plasmid expressing U1 snRNA under the control of its native promoter for 24 hours. Total U1 levels in total RNA were then determined by RT-qPCR. The RNA input was normalized to 5S rRNA. Data are represented as mean + SD. Quantitation of U1 snRNA assembled into snRNPs immunopurified with anti-Sm antibodies, showed the same U1 increase (data not shown).

**Supplementary Fig 3.**
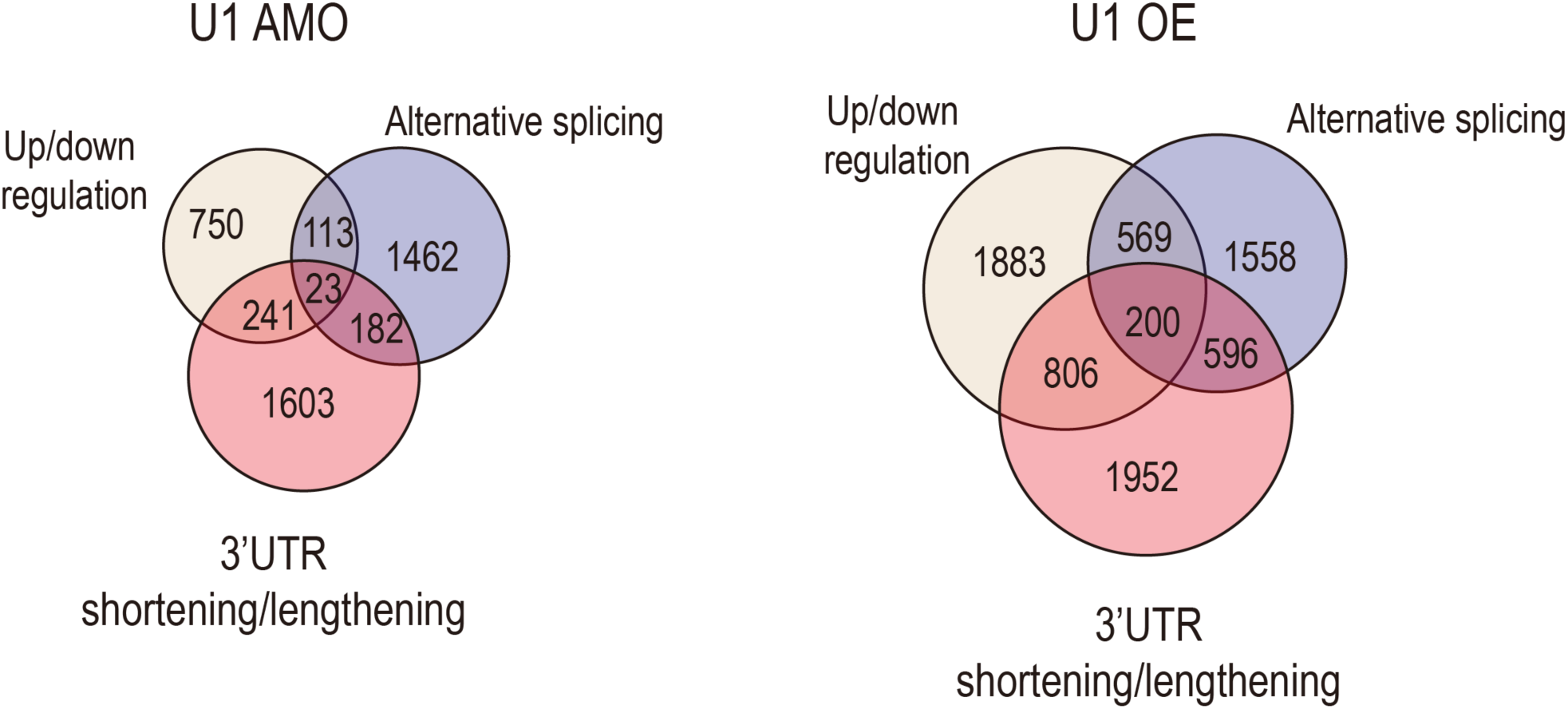
Transcriptome changes induced by U1 AMO and U1 over-expression. Venn diagrams showed the number of overlapped genes from 3 different transcriptome changes: up- and down-regulation, 3’UTR length change, and AS in U1 AMO and U1 OE samples.

**Supplementary Fig 4.**
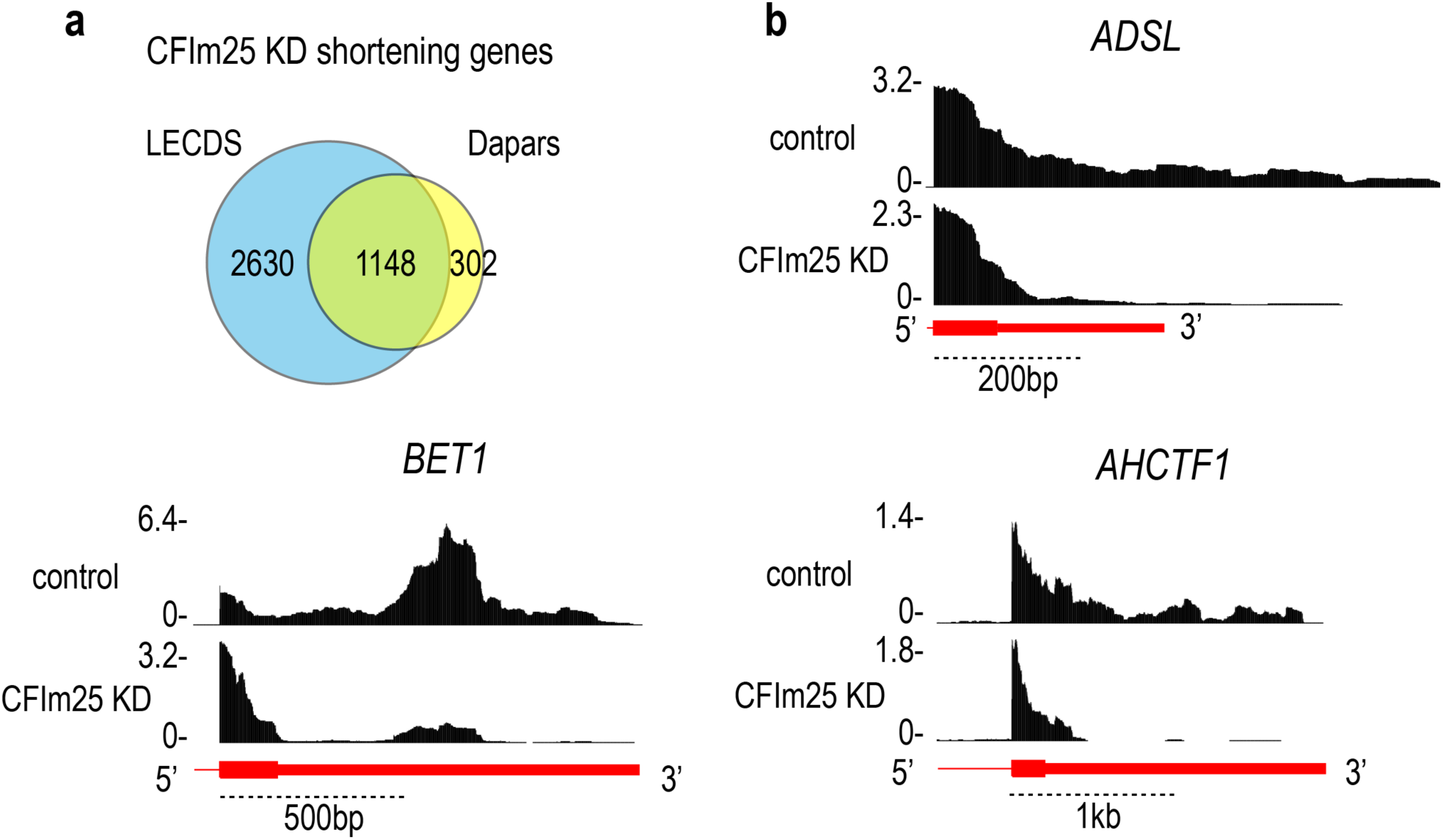
Validation of LECDS calculation. **a** Significant overlap (79%) between 3’UTR shortening genes captured by LECDS and DaPars^14^ calculations. **b** Genome browser examples show the 3’UTR shortening genes captured by LECDS but not by DaPars.

**Supplementary Fig 5.**
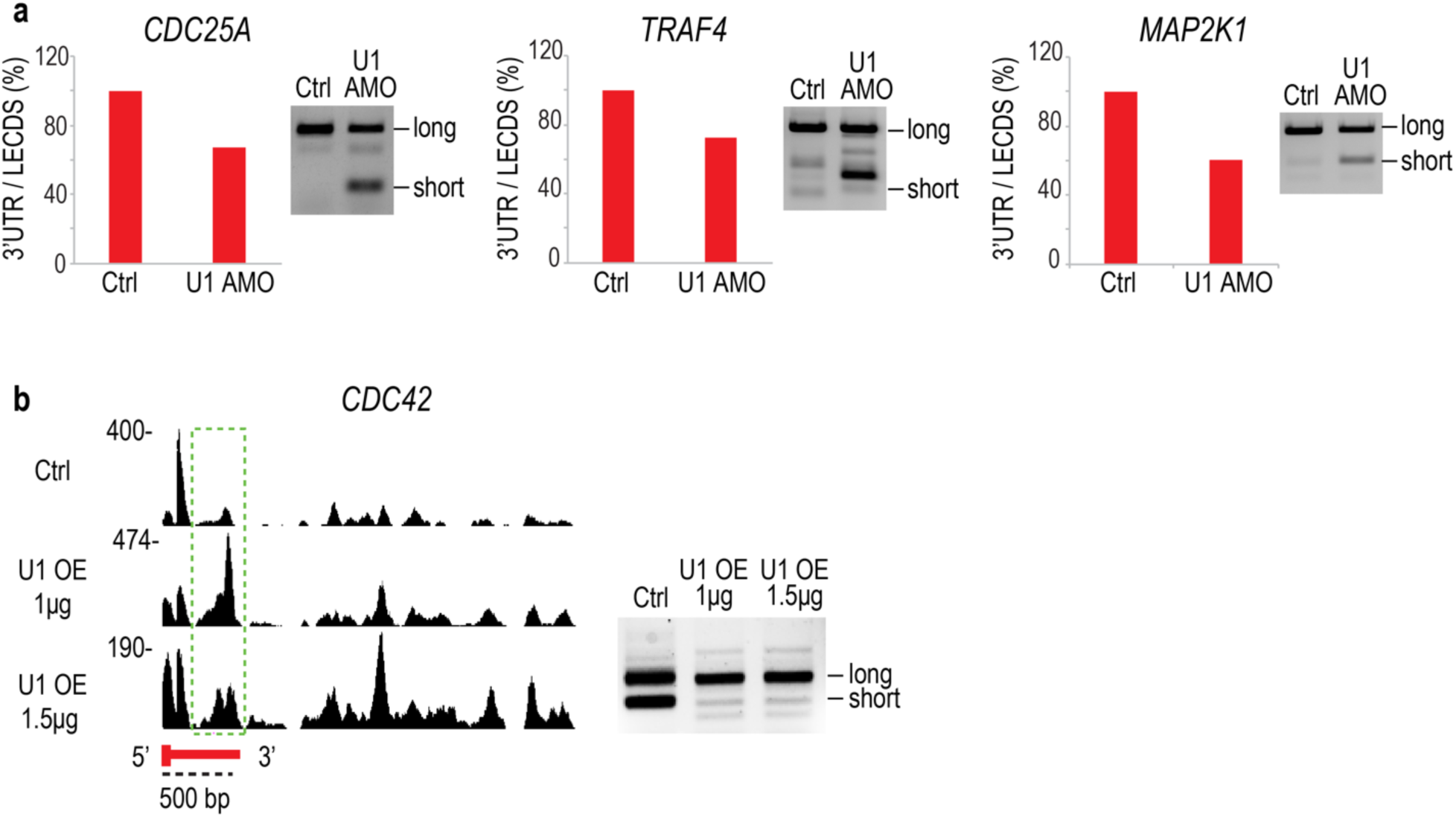
Validation of 3’ UTR length changes after U1 AMO and U1 over-expression. Cells were transfected with control or U1 AMO 62.5 pmole for 8 hours, or U1 expression plasmid for 24 hours. Total RNA was extracted using Trizol followed by selection of nascent transcripts. **a** Histogram shows a percentage of 3’UTR / LECDS in each shortening gene that was selected by LECDS, and which was confirmed by 3’ RACE. **b** Green dotted boxes in *CDC42*’s 3’UTR indicate the region of the 3’UTRs that was affected by U1 OE, and which was confirmed by 3’ RACE.

**Supplementary Fig 6.**
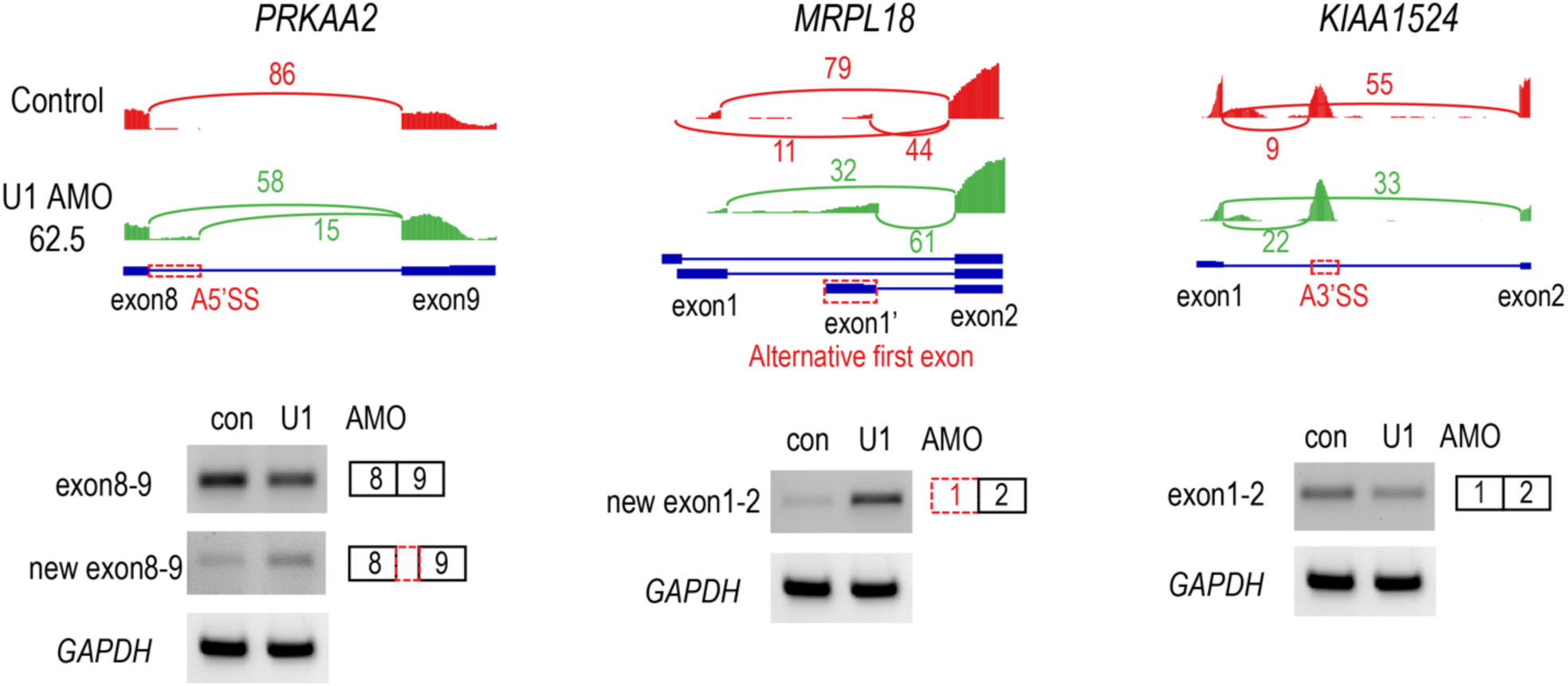
RT-PCR validation of U1 AMO induces alternative splicing changes. HeLa cells transfected with control or U1 AMO 62.5 pmole were incubated with 4-shU in last 2 hours at 6 hours post transfection. 4-shU labeled RNA were purified and used for RT-PCR to confirm the alternative splicing change in each example gene.

**Supplementary Fig 7.**
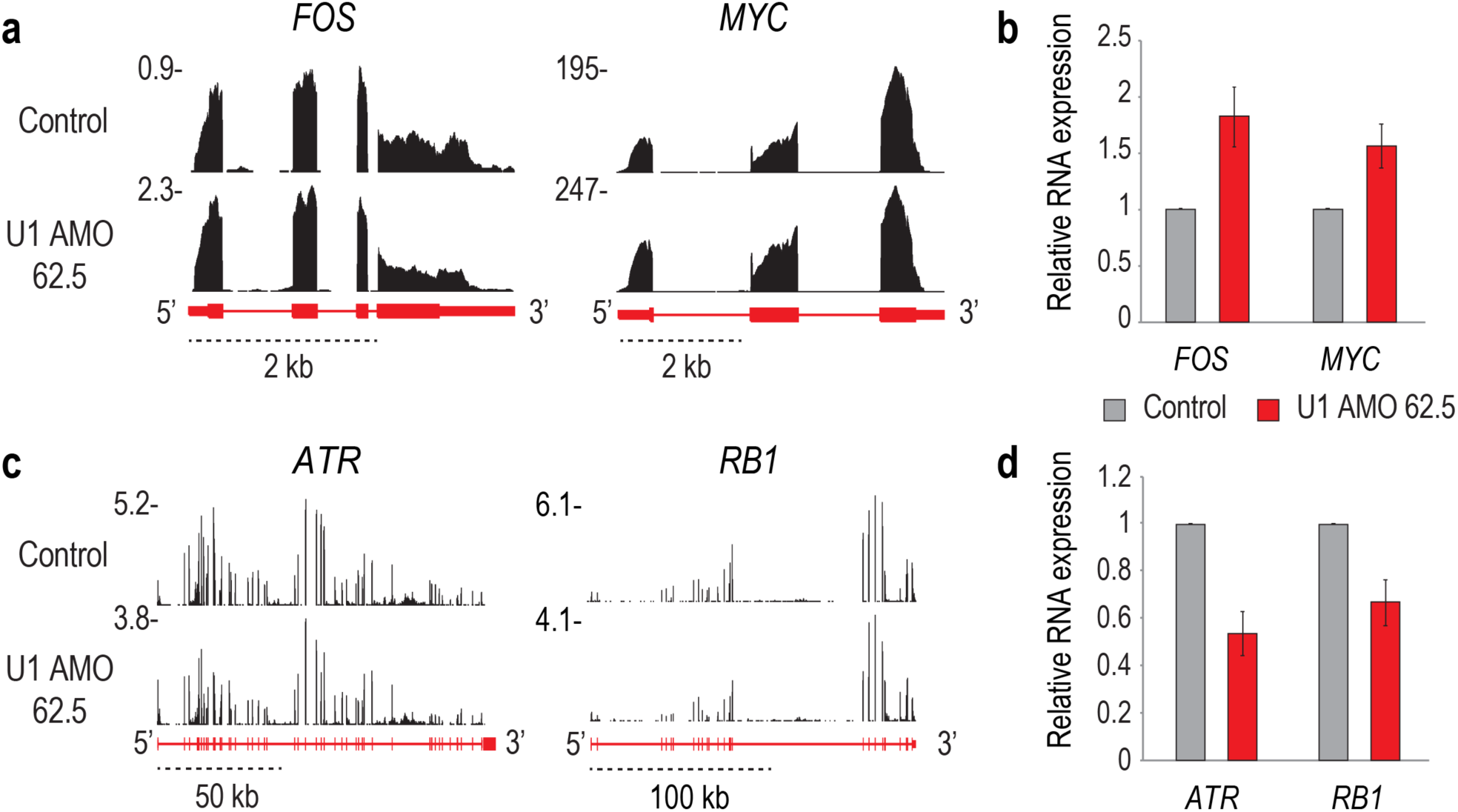
Validation of increased and decreased genes in low U1 AMO. Oncogenes and cancer related genes are increased and genes related to cell cycle and tumor suppressor function are decreased in the low U1 AMO. Genome browser view shows example of up-regulated genes, *FOS* and *MYC* (**a**), and down-regulated genes, *ATR* and *RB1* (**c**). Histogram shows up-regulated genes (**b**), and down-regulated genes (**d**) analyzed by RT-qPCR (mean ± SD; n=3, independent cell cultures). HeLa cells were incubated with 4-shU for last 2 hours after 6 hours of transfection with control or U1 AMO 62.5 pmole. 4-shU labeled RNA were extracted then ERCC RNA spike-in controls added to normalize the RT-qPCR data^7^.

**Supplementary Fig 8.**
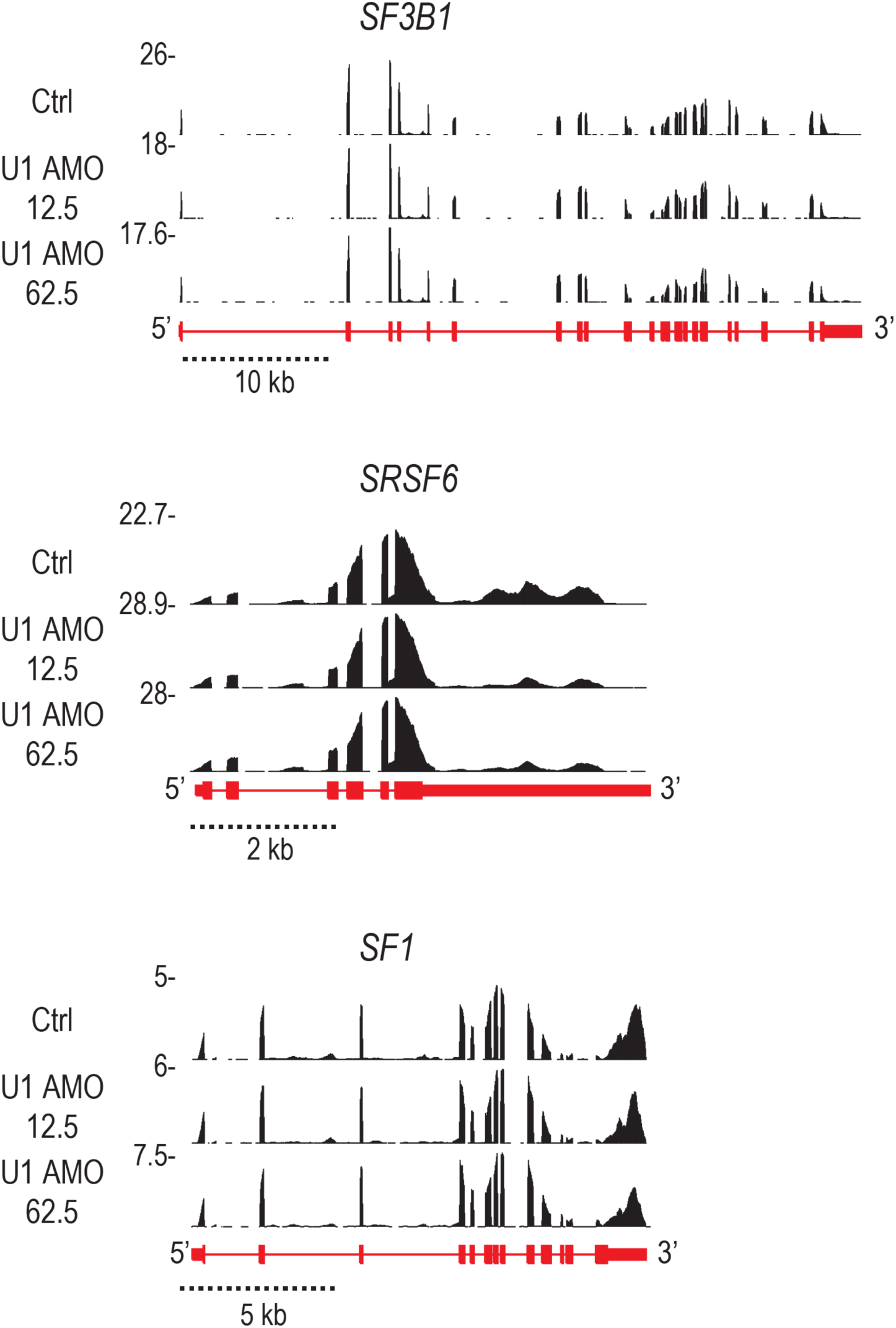
U1 AMO induces expression level changes in splicing factors. Genome browser view shows *SF3B1* decrease and *SRSF6* and *SF1* increases in both U1 AMO 12.5 and 62.5.

**Supplementary Fig 9.**
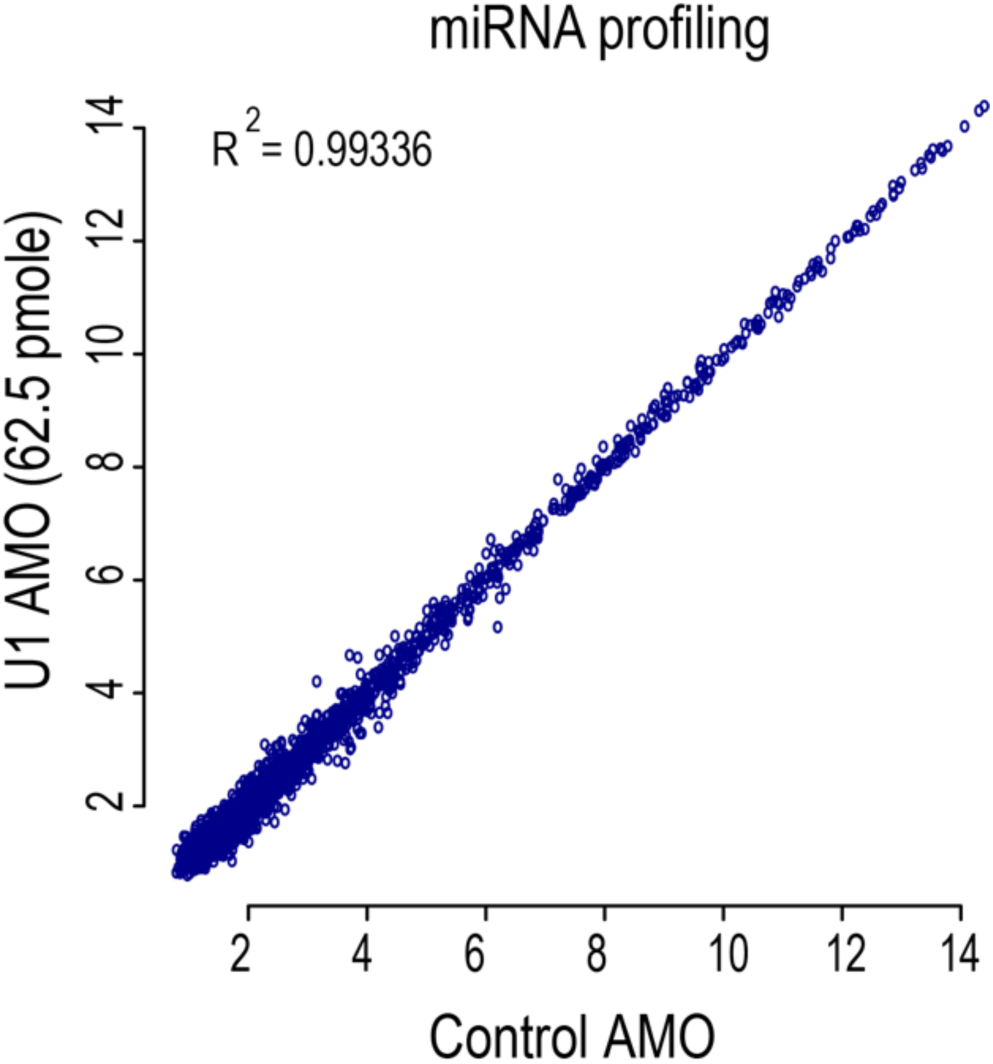
Moderate U1 decrease doesn’t affect steady state miRNA level. A scatter plot of miRNAs expression levels in control cells versus cells transfected with moderate U1 AMO (62.5 pmole) for 8 hours. RNA extraction, microarray used, and analysis are as described in Methods.

**Supplementary Fig 10.**
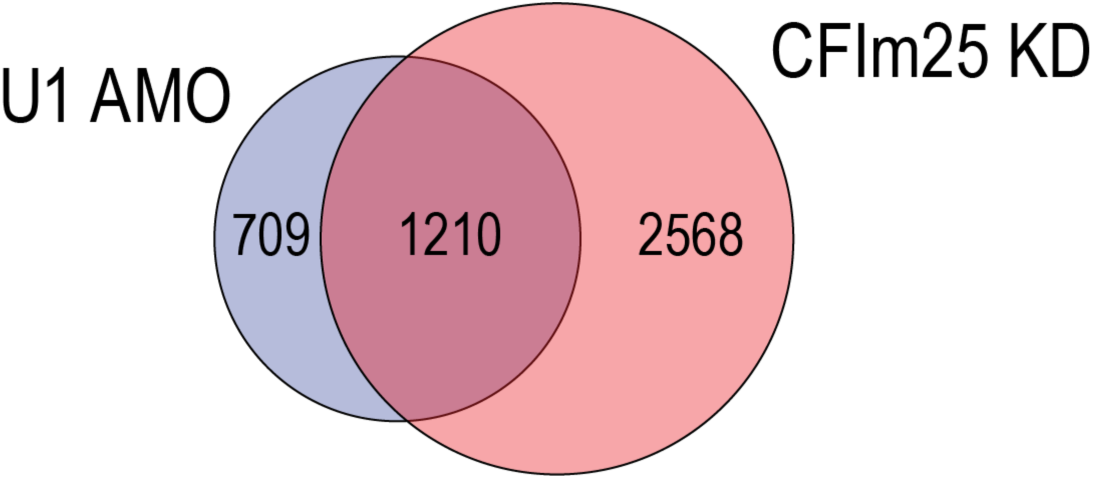
Both low U1 AMO and CFIm25/CPSF5 knockdown cause widespread 3’UTR shortening. Venn diagram of LECDS identified genes with 3’UTR shortening in both 12.5 and 62.5 pmole U1 AMO treatment (blue; 1,919 genes) versus CFIm25/CPSF5 knockdown (red; 3,778 genes) in HeLa cells.

## Supplementary Tables

**Supplementary Table 1.**
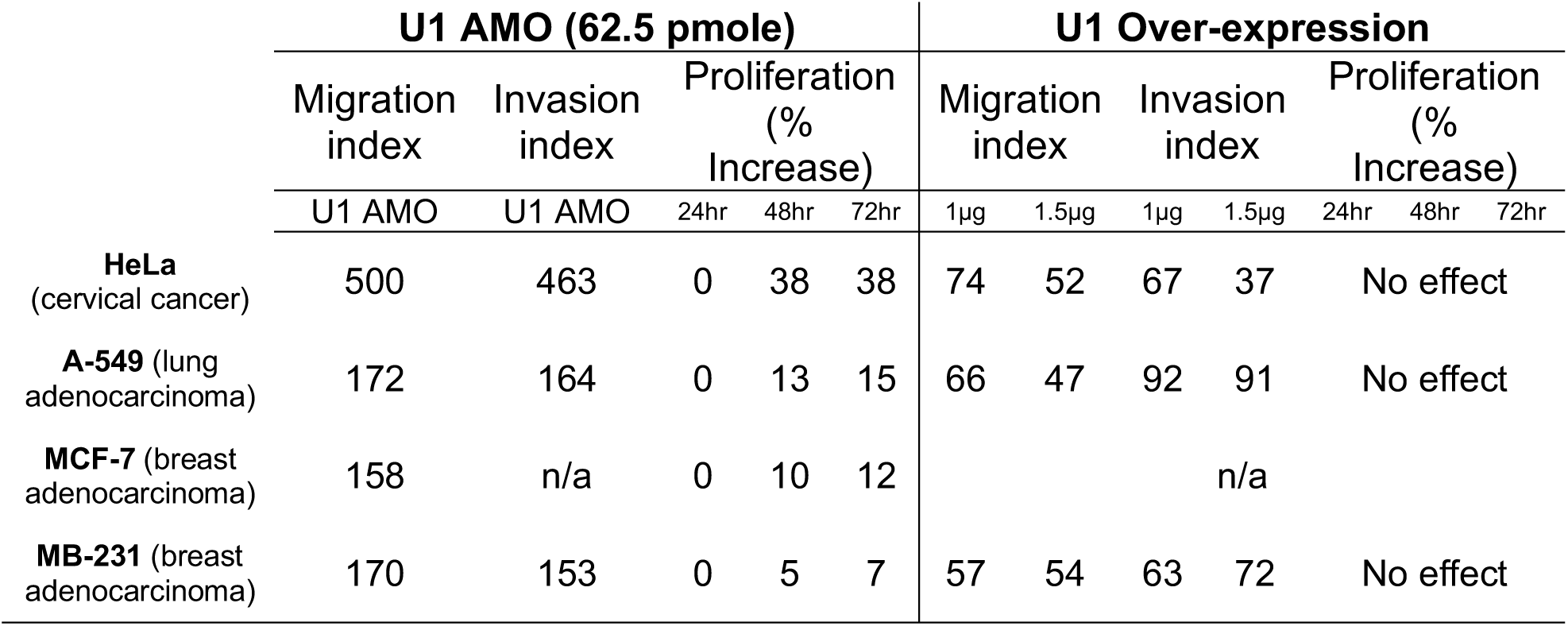
U1 level modulates migration and invasion in cancer cell lines. Migration and invasion changes with U1 AMO or U1 OE were determined as described in Fig 1 on HeLa, A-549, MCF-7, and MB-231 cells. Migration and invasion indexes are presented as percent of control for cells treated with U1 AMO (62.5 pmole) or U1 OE (both 1µg and 1.5µg) for 24 hours. Cell proliferation was assayed daily over three days.

**Supplementary Table 2.** Summary of RNA-seq Data. Total number of reads, mapped reads and the percentage of mapped reads for each RNA-seq sample are presented.

|  | U1 AMO (pmole) |  |  | U1 Over-expression |  |  |
| --- | --- | --- | --- | --- | --- | --- |
| Reads | Control AMO | 12.5 | 62.5 | Empty vector | 1µg | 1.5µg |
| Total | 64,965,053 | 57,555,134 | 60,562,764 | 226,028,904 | 185,947,498 | 207,313,315 |
| Mapped | 47,916,977 | 41,016,817 | 42,980,600 | 136,133,739 | 92,804,677 | 123,786,737 |
| % Mapped | 73.80% | 71.30% | 71.00% | 60.20% | 49.90% | 59.70% |

**Supplementary Table 3.** Transcriptome changes induced by U1 level modulation with U1 AMO or U1 over-expression in HeLa cells compared to controls. Shown are the numbers of genes affected at each dose. mRNA up-regulated and down-regulated genes were determined using GFold^53^ differential expression software with the following cutoffs: fold change >1.5 fold and p-value < 0.01. 3’UTR length changes were determined by calculating the ratio of RNA-seq reads in the last exon over the coding region in the last exon, abbreviated in the text as LECDS, in the indicated conditions versus their controls. Splicing changes were analyzed by JUM^15^ with the following parameters: |ΔΨ|≥ 10%, p-value < 0.05. Abbreviations for alternative splicing events: Alternative 5’/3’ Splice Sites (A5’SS, A3’SS), Cassette Exon (CE), Intron Retention (IR), Mutually Exclusive Exons (MXE).

|  |  | U1 AMO (pmole) |  |  | U1 Over-expression |  |  |
| --- | --- | --- | --- | --- | --- | --- | --- |
| Transcriptome Changes |  | 12.5 | 62.5 | overlap | 1µg | 1.5µg | overlap |
| Differential expression | Up-regulated | 525 | 787 | 309 (59%) | 1249 | 1594 | 1107 (89%) |
|  | Down-regulated | 969 | 1402 | 818 (84%) | 2418 | 4430 | 2351 (97%) |
| 3'UTR length | Total shortened | 2530 | 2630 | 1919 (76%) | 563 | 1063 | 344 (61%) |
|  | Total lengthened | 324 | 218 | 130 (60%) | 4615 | 3831 | 3210 (84%) |
| Alternative Splicing | A5'SS | 1542 | 1547 | 702 (45%) | 1775 | 1653 | 817 (49%) |
|  | A3'SS | 1190 | 1163 | 499 (43%) | 1508 | 1380 | 647 (47%) |
|  | CE | 721 | 705 | 247 (35%) | 820 | 725 | 275 (38%) |
|  | IR | 334 | 483 | 152 (31%) | 1859 | 1769 | 1059 (60%) |
|  | MXE | 6 | 7 | 3 (43%) | 4 | 7 | 3 (43%) |

**Supplementary Table 4.**
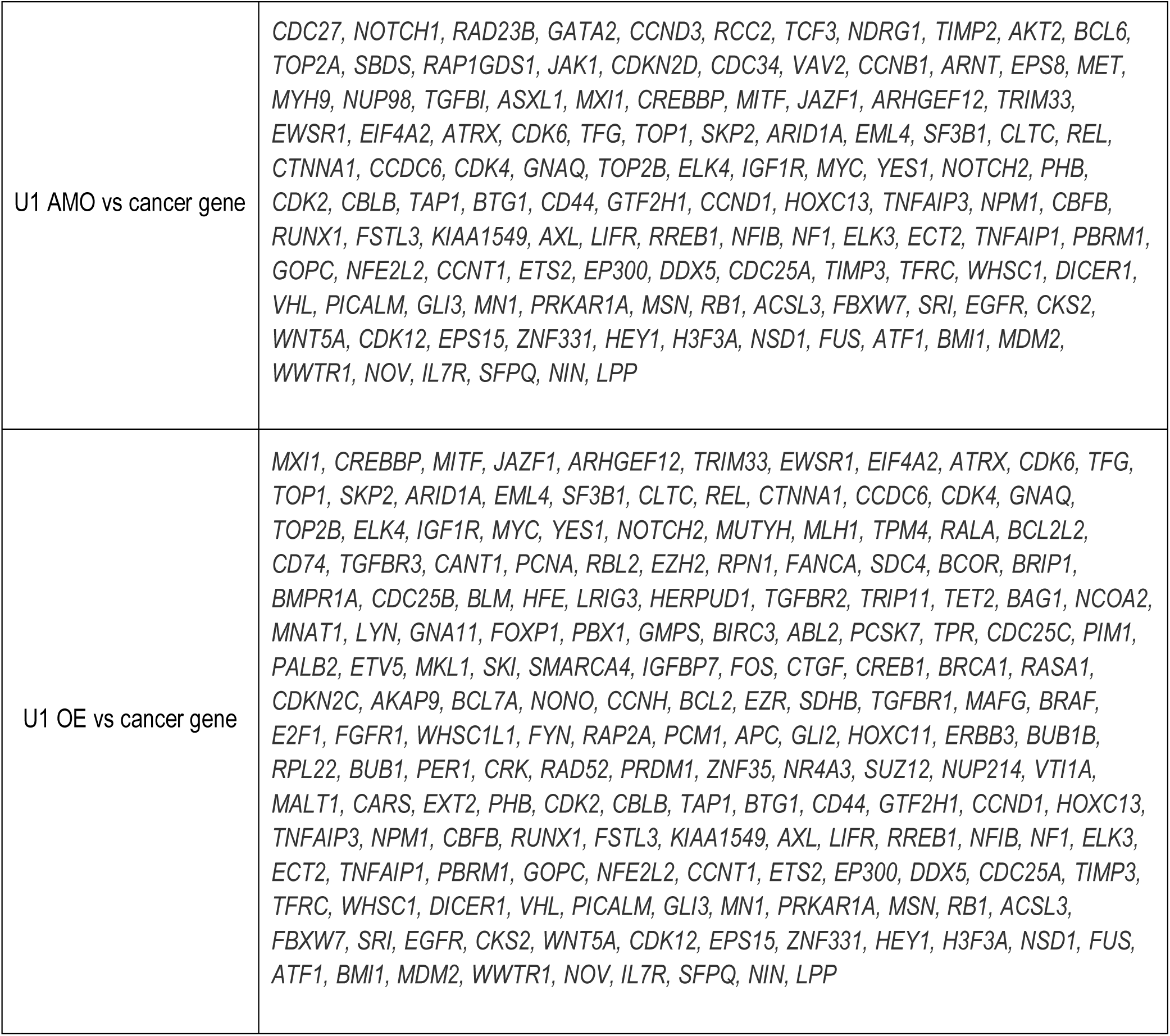
Cancer genes affected by U1 AMO and U1 over-expression. Table showed the overlapped genes between cancer genes and 3’UTR length changed genes in U1 AMO and U1 OE samples.

**Supplementary Table 5.**
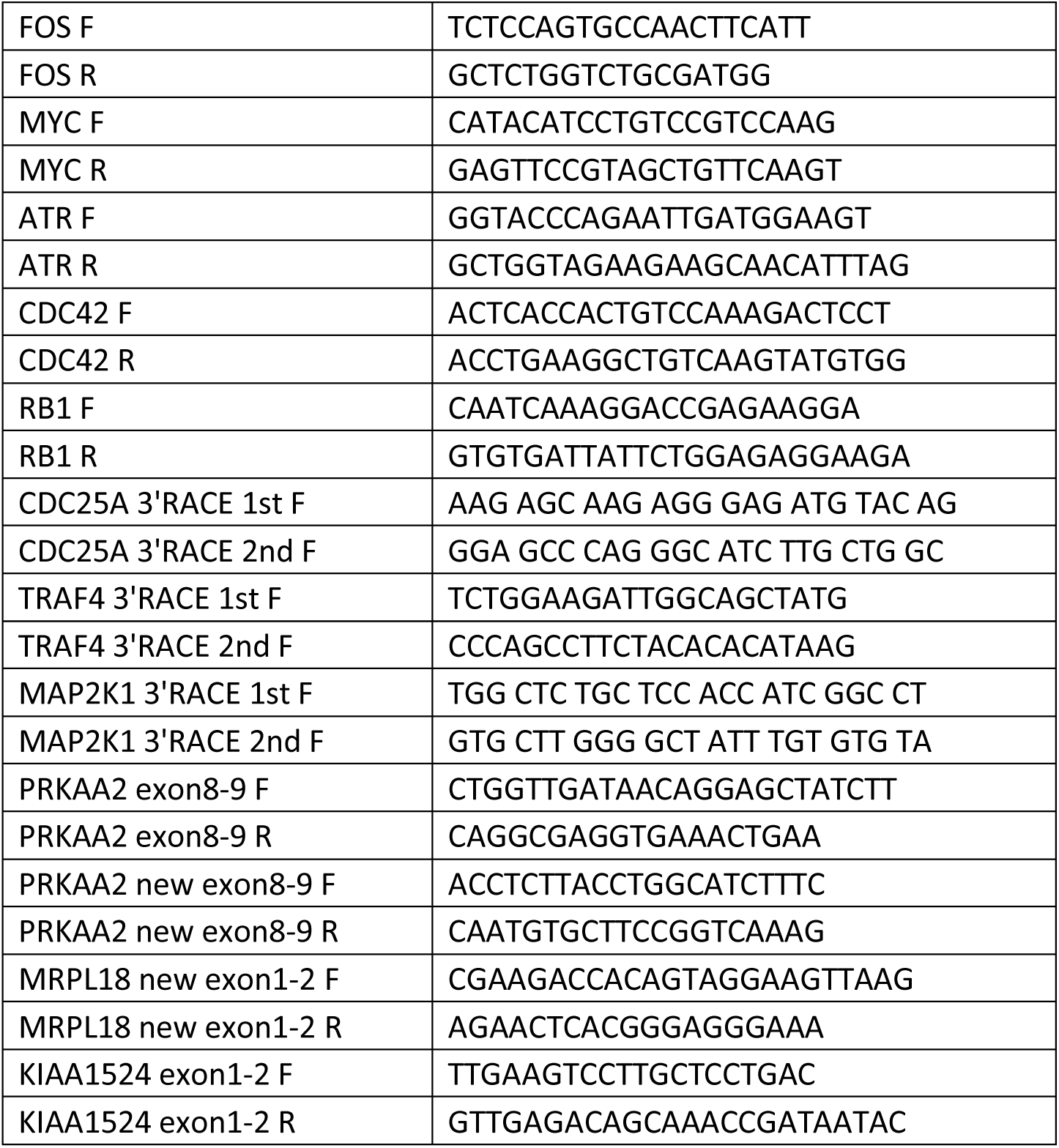
Primers used for RT-qPCR, 3’RACE and RT-PCR (Supplementary Figs 5-7)

## References

1. Flavell, S.W. et al. Genome-wide analysis of MEF2 transcriptional program reveals synaptic target genes and neuronal activity-dependent polyadenylation site selection. Neuron 60, 1022–38 (2008).

2. Sandberg, R., Neilson, J.R., Sarma, A., Sharp, P.A. & Burge, C.B. Proliferating cells express mRNAs with shortened 3’ untranslated regions and fewer microRNA target sites. Science 320, 1643–7 (2008).

3. Ji, Z. & Tian, B. Reprogramming of 3’ untranslated regions of mRNAs by alternative polyadenylation in generation of pluripotent stem cells from different cell types. PLoS One 4, e8419 (2009).

4. Mayr, C. & Bartel, D.P. Widespread shortening of 3’UTRs by alternative cleavage and polyadenylation activates oncogenes in cancer cells. Cell 138, 673–84 (2009).

5. Singh, I. et al. Widespread intronic polyadenylation diversifies immune cell transcriptomes. Nat Commun 9, 1716 (2018).

6. Lee, S.H. et al. Widespread intronic polyadenylation inactivates tumour suppressor genes in leukaemia. Nature 561, 127–131 (2018).

7. Oh, J.M. et al. U1 snRNP telescripting regulates a size-function-stratified human genome. Nat Struct Mol Biol 24, 993–999 (2017).

8. Kaida, D. et al. U1 snRNP protects pre-mRNAs from premature cleavage and polyadenylation. Nature 468, 664–8 (2010).

9. Berg, M.G. et al. U1 snRNP determines mRNA length and regulates isoform expression. Cell 150, 53–64 (2012).

10. Matera, A.G. & Wang, Z. A day in the life of the spliceosome. Nat Rev Mol Cell Biol 15, 108–21 (2014).

11. Niibori, Y., Hayashi, F., Hirai, K., Matsui, M. & Inokuchi, K. Alternative poly(A) site-selection regulates the production of alternatively spliced vesl-1/homer1 isoforms that encode postsynaptic scaffolding proteins. Neurosci Res 57, 399–410 (2007).

12. Matter, N. & Konig, H. Targeted ‘knockdown’ of spliceosome function in mammalian cells. Nucleic Acids Res 33, e41 (2005).

13. Li, W. et al. Systematic profiling of poly(A)+ transcripts modulated by core 3’ end processing and splicing factors reveals regulatory rules of alternative cleavage and polyadenylation. PLoS Genet 11, e1005166 (2015).

14. Masamha, C.P. et al. CFIm25 links alternative polyadenylation to glioblastoma tumour suppression. Nature 510, 412–6 (2014).

15. Wang, Q. & Rio, D.C. JUM is a computational method for comprehensive annotation-free analysis of alternative pre-mRNA splicing patterns. Proc Natl Acad Sci U S A 115, E8181–E8190 (2018).

16. Gilad, S. et al. Genotype-phenotype relationships in ataxia-telangiectasia and variants. Am J Hum Genet 62, 551–61 (1998).

17. Yoshida, K. et al. Frequent pathway mutations of splicing machinery in myelodysplasia. Nature 478, 64–9 (2011).

18. Ilagan, J.O. et al. U2AF1 mutations alter splice site recognition in hematological malignancies. Genome Res 25, 14–26 (2015).

19. Kim, E. et al. SRSF2 Mutations Contribute to Myelodysplasia by Mutant-Specific Effects on Exon Recognition. Cancer Cell 27, 617–30 (2015).

20. Alsafadi, S. et al. Cancer-associated SF3B1 mutations affect alternative splicing by promoting alternative branchpoint usage. Nat Commun 7, 10615 (2016).

21. Futreal, P.A. et al. A census of human cancer genes. Nat Rev Cancer 4, 177–83 (2004).

22. Nicoloso, M.S., Spizzo, R., Shimizu, M., Rossi, S. & Calin, G.A. MicroRNAs--the micro steering wheel of tumour metastases. Nat Rev Cancer 9, 293–302 (2009).

23. Nana-Sinkam, S.P. & Croce, C.M. Non-coding RNAs in cancer initiation and progression and as novel biomarkers. Mol Oncol 5, 483–91 (2011).

24. Ebert, M.S. & Sharp, P.A. Roles for MicroRNAs in Conferring Robustness to Biological Processes. Cell 149, 515–24 (2012).

25. Mendell, J.T. & Olson, E.N. MicroRNAs in stress signaling and human disease. Cell 148, 1172–87 (2012).

26. Johnson, C.D. et al. The let-7 microRNA represses cell proliferation pathways in human cells. Cancer Res 67, 7713–22 (2007).

27. Lee, S.O. et al. MicroRNA15a modulates expression of the cell-cycle regulator Cdc25A and affects hepatic cystogenesis in a rat model of polycystic kidney disease. J Clin Invest 118, 3714–24 (2008).

28. Wang, P. et al. microRNA-21 negatively regulates Cdc25A and cell cycle progression in colon cancer cells. Cancer Res 69, 8157–65 (2009).

29. Paranjape, T. et al. A 3’-untranslated region KRAS variant and triple-negative breast cancer: a case-control and genetic analysis. Lancet Oncol 12, 377–86 (2011).

30. Milde-Langosch, K. The Fos family of transcription factors and their role in tumourigenesis. Eur J Cancer 41, 2449–61 (2005).

31. Silvestre, D.C., Gil, G.A., Tomasini, N., Bussolino, D.F. & Caputto, B.L. Growth of peripheral and central nervous system tumors is supported by cytoplasmic c-Fos in humans and mice. PLoS One 5, e9544 (2010).

32. Takagaki, Y., Seipelt, R.L., Peterson, M.L. & Manley, J.L. The polyadenylation factor CstF-64 regulates alternative processing of IgM heavy chain pre-mRNA during B cell differentiation. Cell 87, 941–52 (1996).

33. Di Giammartino, D.C., Nishida, K. & Manley, J.L. Mechanisms and consequences of alternative polyadenylation. Mol Cell 43, 853–66 (2011).

34. de Klerk, E. et al. Poly(A) binding protein nuclear 1 levels affect alternative polyadenylation. Nucleic Acids Res (2012).

35. Martin, G., Gruber, A.R., Keller, W. & Zavolan, M. Genome-wide Analysis of Pre-mRNA 3’ End Processing Reveals a Decisive Role of Human Cleavage Factor I in the Regulation of 3’ UTR Length. Cell Rep 1, 753–63 (2012).

36. Yao, C. et al. Transcriptome-wide analyses of CstF64-RNA interactions in global regulation of mRNA alternative polyadenylation. Proc Natl Acad Sci U S A (2012).

37. Jenal, M. et al. The poly(a)-binding protein nuclear 1 suppresses alternative cleavage and polyadenylation sites. Cell 149, 538–53 (2012).

38. Gruber, A.R., Martin, G., Keller, W. & Zavolan, M. Means to an end: mechanisms of alternative polyadenylation of messenger RNA precursors. Wiley Interdiscip Rev RNA 5, 183–96 (2014).

39. Yu, K. et al. A precisely regulated gene expression cassette potently modulates metastasis and survival in multiple solid cancers. PLoS Genet 4, e1000129 (2008).

40. Park, H.J. et al. 3’ UTR shortening represses tumor-suppressor genes in trans by disrupting ceRNA crosstalk. Nat Genet 50, 783–789 (2018).

41. Cheng, Z. et al. Gene expression profiling reveals U1 snRNA regulates cancer gene expression. Oncotarget 8, 112867–112874 (2017).

42. Hanahan, D. & Weinberg, R.A. Hallmarks of cancer: the next generation. Cell 144, 646–74 (2011).

43. Sherr, C.J. Principles of tumor suppression. Cell 116, 235–46 (2004).

44. Henrich, K.O., Schwab, M. & Westermann, F. 1p36 tumor suppression--a matter of dosage? Cancer Res 72, 6079–88 (2012).

45. Chiu, A.C. et al. Transcriptional Pause Sites Delineate Stable Nucleosome-Associated Premature Polyadenylation Suppressed by U1 snRNP. Mol Cell 69, 648–663 e7 (2018).

46. Almada, A.E., Wu, X., Kriz, A.J., Burge, C.B. & Sharp, P.A. Promoter directionality is controlled by U1 snRNP and polyadenylation signals. Nature 499, 360–3 (2013).

47. Vorlova, S. et al. Induction of antagonistic soluble decoy receptor tyrosine kinases by intronic polyA activation. Mol Cell 43, 927–39 (2011).

48. Langemeier, J. et al. A complex immunodeficiency is based on U1 snRNP-mediated poly(A) site suppression. EMBO J 31, 4035–44 (2012).

49. Tsujii, M. et al. Cyclooxygenase regulates angiogenesis induced by colon cancer cells. Cell 93, 705–16 (1998).

50. Kramer, N. et al. In vitro cell migration and invasion assays. Mutat Res (2012).

51. Dobin, A. et al. STAR: ultrafast universal RNA-seq aligner. Bioinformatics 29, 15–21 (2013).

52. Mortazavi, A., Williams, B.A., McCue, K., Schaeffer, L. & Wold, B. Mapping and quantifying mammalian transcriptomes by RNA-Seq. Nat Methods 5, 621–8 (2008).

53. Feng, J. et al. GFOLD: a generalized fold change for ranking differentially expressed genes from RNA-seq data. Bioinformatics 28, 2782–8 (2012).

